# Non-linear development of EEG coherence in adolescents and young adults shown by the analysis of neurophysiological trajectories and their covariance

**DOI:** 10.1101/2024.03.13.584867

**Authors:** David B. Chorlian, Chella Kamarajan, Jacquelyn L. Meyers, Ashwini K. Pandey, Jian Zhang, Sivan Kinreich, Bernice Porjesz

## Abstract

To contribute to the understanding of changes in the factors governing the development of neural connectivity, the developmental structure of EEG coherence in adolescents and young adults was analyzed using the means, variances, and covariances of high alpha frequency band coherence measures from a set of 27 coherence pairs obtained from a sample of 1426 participants from the COGA study with 5006 observations over ages 12 through 31. Means and covariances were calculated at 96 age centers by a LOESS method.

In the current study, trajectories of covariance matrices considered as individual units were determined by tensorial analysis: calculation of Riemannian geodesic (non-Euclidean) distances between matrices and application of both linear and non-linear dimension reduction techniques to these distances. Results were evaluated by bootstrap methods.

Mean coherence trajectories for males and females were very similar, showing a steady upward trend from ages 12 to 20 which diminishes gradually from 20 to 25 and reaches stability from 25 to 31. In contrast, the individual covariance trajectories of males and female differed, with the male covariance levels becoming greater than that of females during the developmental process. Tensorial determination of the distances from the initial covariance matrix of subsequent covariance matrices to age 20 had the same trajectory as the mean coherence values. Tensorial determination of the trajectories of the covariance matrices of males and females based on their all pairs geodesic distances revealed a non-linear pattern in the multi-dimensional space of each of the trajectories: A steady increase in one dimension is accompanied by deviations from it peaking at age 20 which have both transient and lasting effects. There is a precise temporal parallelism of this pattern of covariance in males and females, while there is a consistent distance between the male and female covariance structures throughout the developmental process. Between region differences (anterior-posterior) within each sex are greater than between sex differences within regions. Examining development using multiple methods provides unique insight into the developmental process.

## 1 Introduction

In this study of the developmental trajectories of EEG coherence, the age and region specific covariance between coherence measures is studied as an indication of the age and sex varying topographic differences between the factors influencing the development of neural connectivity, as a necessary complement to the study of the means and variances of the individual coherence measures. Examination of the trajectories of coherence by themselves cannot determine whether parallel trajectories are the result of a common underlying process or several independent but parallel processes.

It must be emphasized that the analysis of the relations between covariance matrices as presented in this study provides no direct information about which elements of the matrices, i.e. which specific sets of coherence pairs, account for the developmental features obtained from the analysis. Topographical specificity can only be obtained by analysis from topographically specific submatrices or from the individual covariance trajectories themselves. A subsequent study will undertake that project in the framework provided by the results outlined below.

The guiding ideas of this study are well expressed by Mitteroecker and Bookstein (2009):

> Whenever development varies among individuals, the population covariance matrix is subject to ontogenetic change, … When new processes emerge in the course of development (by way of, for example, expression of new genes, new epigenetic interactions, or environmental influences), a novel pattern of variances and covariances is superimposed on top of the previous population covariance structure … Similarly, when developmental processes cease at some stage, they stop contributing to the covariance structure of a population. The timing of cessation and initiation of factors may vary itself among individuals and contribute to the observed phenotypic variation. A careful statistical tracking of age or stage-specific covariance matrices over ontogeny hence might permit inferences about the timing and phenotypic effects of developmental processes.

## 2 Materials and Methods

### 2.1 Sample

As part of the Collaborative Study on the Genetics of Alcoholism (COGA) (Begleiter et al., 1995; Agrawal et al., 2023), EEG data were collected from adolescent and young adult offspring of members of initially recruited subjects from families densely affected with AUD as well as community comparison families. In this study EEG coherence measures in the high alpha band (9-12 Hz.) from 27 coherence pairs were obtained from a sample of 1426 participants, children of the originally recruited sample, with 5006 assessments from ages 12 through 31 were included in the analysis. The sample and coherence measures were chosen to be those used in our recent papers on the genetic association of EEG coherence values and various PGS measures of psychiatric conditions (Meyers et al., 2019, 2021), partially to advance the understanding of the phenotypic characteristics of the measures analyzed in those works.

### 2.2 Data Collection and Processing

Data was collected and coherence was calculated as described in Chorlian et al. (2009) and Meyers et al. (2019). Briefly, eyes-closed resting-state EEG data was recorded from the electrodes of the 10-20 system for a period of 256 seconds, bipolar channels were created by subtraction between sagittally and laterally located pairs of electrodes, and coherence was calculated between 27 pairings of similarly oriented bipolar channels (21 between sagittal channels and 6 between lateral channels) by a Fourier transform method. Two complementary subsets of 10 members each of the between sagittal coherence pairs, anterior (coherence between frontal-central bipolar channels) and posterior (coherence between central-parietal bipolar channels) with identical topographic structure (see Figure 1), were subject to the same analysis as the complete system. The sagittal pairs contain both interhemispheric and intrahemispheric pairs. The 6 lateral coherence coherence pairs are intrahemispheric pairs with left-right symmetry (See Figure 2). A complete table is provided in Section 6.1.

**Figure 1:**
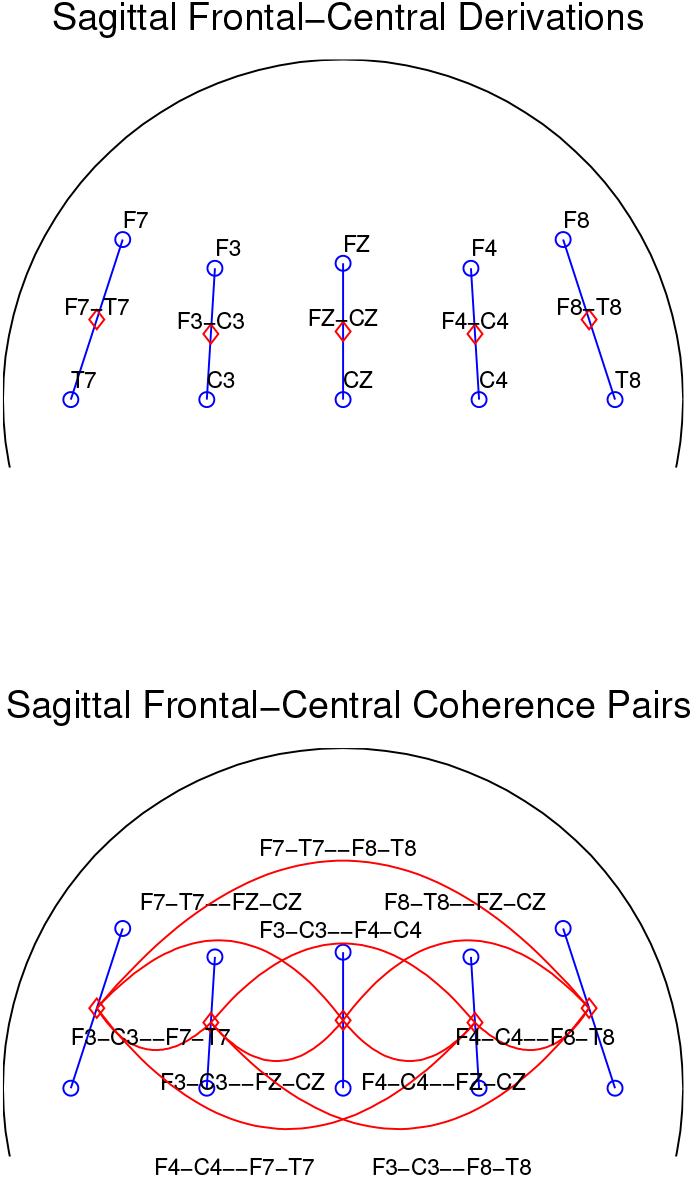
Upper panel: Frontal-central electrodes and sagittal bipolar derivations. Blue lines connect the electrodes forming the bipolar channels. Lower panel: Blue lines connect the electrodes forming the bipolar channels. Red lines connect midpoints of coherence pairs for all pairs calculations. Central-parietal is similar.

**Figure 2:**
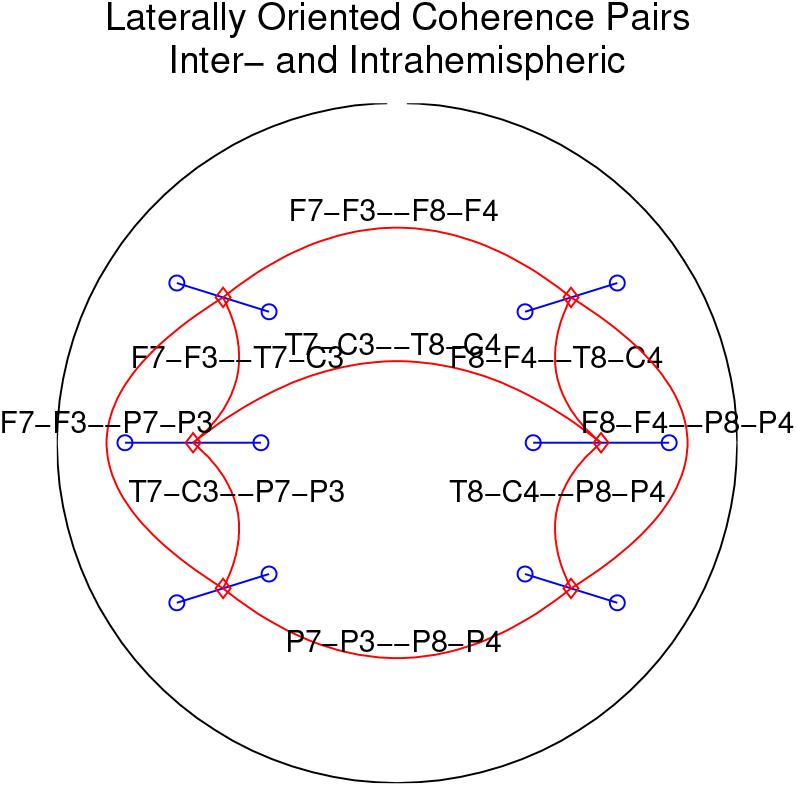
Derivations and coherence pairs for lateral intra- and interhemispheric coherence pairs. Interhemispheric pairs are not used in this analysis.

**Figure 3:**
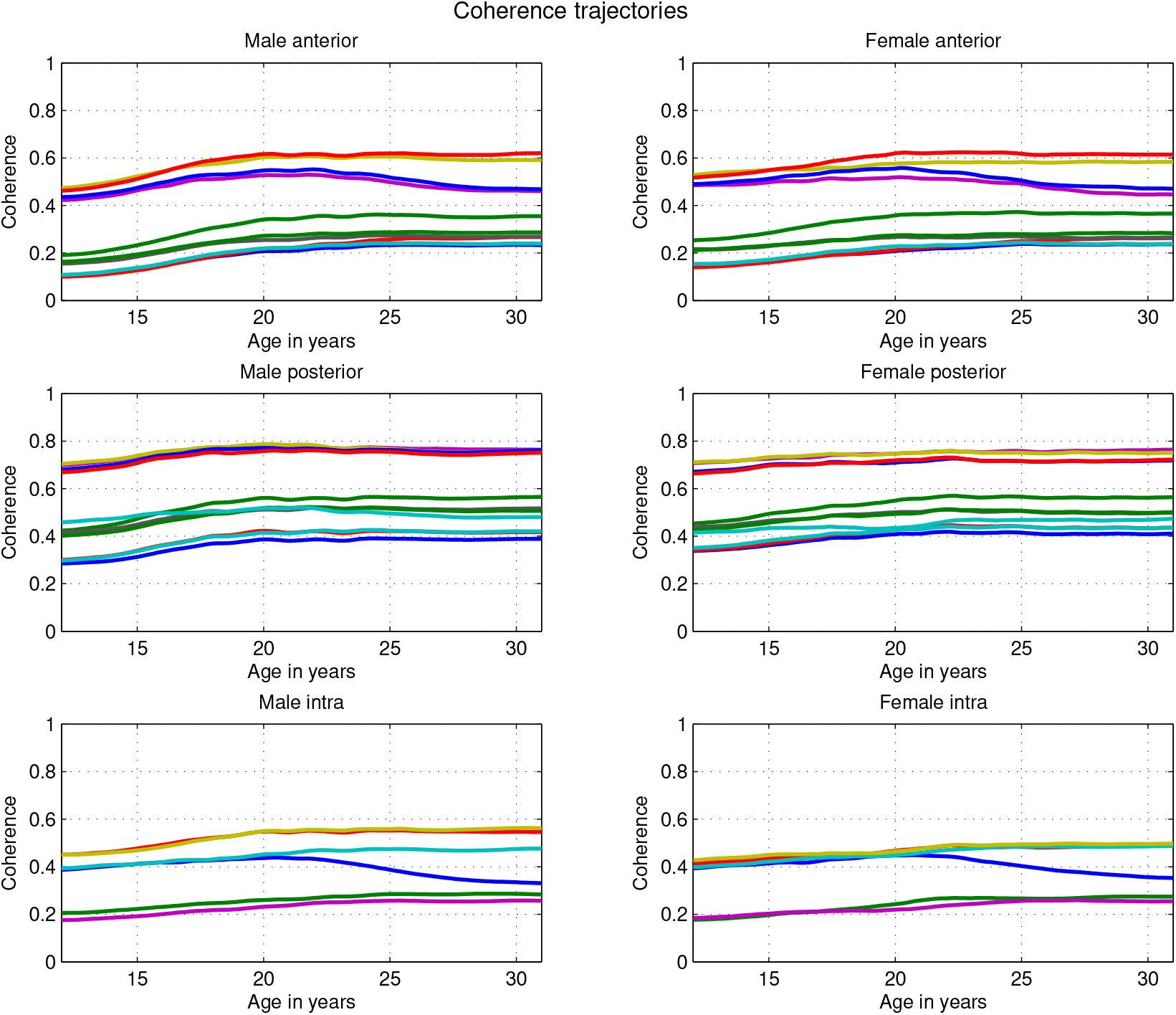
High Alpha band trajectories of the 27 bipolar coherence pairs used in this study. Note that there is little difference between male and female trajectories.

### 2.3 Methods of Analysis

The initial phase of the analysis provides the means, variances, covariances, and correlations of the coherence values of the data set. Although seven frequency bands were available for analysis, only high alpha was analyzed based on its major occurrence in resting state EEG and its use in our previous work (Meyers et al., 2019, 2021). Values at 96 age centers from ages 12 to 31 for each sex were calculated by a local linear regression (LOESS) method and and an adaptation of that method to calculate covariances initially described in Chorlian et al. (2015). Briefly, the weights used in the regression calculation were employed in a weighted covariance analysis. The bandwidth for the kernel (Epanechnikov) was taken as 0.6 as in Chorlian et al. (2015). Quartiles of the values of covariance and correlations matrices were also calculated.

Males and females are analyzed separately in order to properly estimate sex differences in developmental trajectories, since previous work (Cousminer et al., 2013; Chorlian et al., 2015, 2017; Meyers et al., 2019, 2021; Chorlian et al., 2023) suggests that there may be important differences between male and female developmental patterns, both in terms of specific measures and genetic associations with those measures.

#### 2.3.1 Distances between Covariance Matrices

After the calculation of the trajectories of the individual measures and the trajectories of pairwise correlations/covariances of the individual measures are obtained, the developmental analysis of the structural features of the covariance matrices of the observed measures is performed by the calculation of Riemannian (geodesic) distances between covariance matrices. Covariance matrices are properly analyzed as tensors in the non-Euclidean space of positive definite symmetric (PDS) matrices. The methods of tensorial analysis enable the analysis of the developmental process of a system characterized by a multivariate data set as a single system, rather than simply as a collection of individual items. Tensorial analysis provides only measures of the structural relations between matrices; it does not indicate which elements of the matrices are responsible for the distances calculated.

Geodesic distances between pairs of covariance matrices and geodesics between covariance matrices were calculated by the log-Euclidean method (Arsigny et al., 2006; Fletcher et al., 2004; Dryden et al., 2009) and implemented in Matlab. (See Section 6.2 for the mathematical details.). The pairwise distance calculation was applied to all pairs across age of the sex specific covariance matrices of the entire 27 coherence pairs and to the two regional blocks of coherence pairs (10 pairs each), anterior and posterior, which had identical topographic structure by the 10-20 system of scalp coordinates. This made possible comparisons between sexes and regions. Since there are 96 age specific covariance matrices for each block, there are 4560 pairwise distances calculate for each block. Additionally, for each age span distances between the covariance matrices of corresponding blocks were calculated. This provides a direct measure of the trajectories of between block differences, which could either be between sex distances for the same region or between region distances for the same sex.

In order to evaluate the range of variation within the observed covariance matrices, a permutation method was used in which each of the observed sequence of 96 covariance matrices for males and separately for females had its rows and columns permuted by the same random permutation and the maximum of the pairwise geodesic distances between the corresponding observed and permuted matrices recorded for each of 1000 permutations. The range of values for the maxima was between 9 and 10 for each sex, providing a estimate for the diameter of the set of comparable covariance matrices, in the sense of the within matrix distribution of the covariances.

#### 2.3.2 Trajectories of Covariance Matrices

A trajectory is a spatial representation of a time series of states determined either by a value for each state or a procedure by which the pairwise distances between the states is used to create a location in some space for each state. In this study, the states are the covariance matrices at different age centers, and the distances are the Riemannian (geodesic) distances in the non-Euclidean space of positive definite symmetric (PDS) matrices, as described above. Two different approaches are then used to describe the trajectories as determined by the pairwise distances between the matrices. The first uses multidimensional scaling (MDS) applied to the set of geodesic distances between all pairs of the covariances matrices of each type to provide a Euclidean approximation suitable for large scale description and analysis. The second works directly in the Riemannian space to characterize the progression and deviation of the observed trajectory from a trajectory along a geodesic determined by two matrices on the trajectory, providing a measure of the “non-linearity” of the observed trajectory. The analyses based on direct calculations in the non-Euclidean space of the covariance matrices are based on ideas in Fletcher et al. (2004) and Fletcher and Joshi (2007).

While the Euclidean method is hypothesis free, the Riemannian framework used here is that the trajectories are to be described in terms of a linear geodesic progression from an initial stage to a final stage, with possible excursions from this linear progression extending over different age ranges and in different directions. This framework is chosen because of the mathematical convenience to start with a linear progression, particularly as it makes comparison between different measures more comprehensible if they are placed in a common framework. It should be emphasized that this is a purely exploratory data analysis; no particular hypotheses are advanced about the structure or content of the developmental process. All statistical analysis is post hoc, devoted to the further understanding of features observed in the initial phase of the analysis.

1. Euclidean Method: Multidimensional scaling Multidimensional scaling operates on a set of pairwise distances between objects, and constructs a system of coordinates for a space which “best” represents their distance according one of a number of criteria. Classical multidimensional scaling was used in this study, which provides coordinates for a Euclidean space which best represents the pairwise distances in a least squares sense. Trajectories were determined by use of the classical multidimensional scaling algorithm implemented in Matlab. It should be noted that the signs of the coordinates are arbitrary, in the manner that maps conventionally have north at their upper edge, but the same map displayed upside down would be an equivalent representation of the same area. Visualization and interpretation of the results are provided by plotting the trajectories in each dimension as trajectories along the age axis, rather than a space curve in two or more dimensions. Since the matrices lie in a non-Euclidean space the coordinates generated by MDS will have limited accuracy in enabling the reproduction of the pairwise distances. Just as in PCA, the coordinates are determined in order of the amount of variance explained by the distance along the coordinates. Examination of the eigenvalues of the coordinates showed that the first three coordinates accounted for over 75% of the variance of the trajectories, so only the first three coordinates were used in the analysis. There is no direct way to determine what particular relations between 27 coherence pairs is represented by a particular coordinate, unlike PCA. The principal items for study are the comparison of these trajectories between sexes and regions.
2. Riemannian Method: Geodesic Linearity Analysis Geodesic linearity analysis (GLA) was developed as a simplification of the methods described in Fletcher et al. (2004) and Fletcher and Joshi (2007). Geodesic linearity analysis works directly in the non-Euclidean space of the covariances. In this method, the geodesic between an initial matrix and a final matrix was determined, and then it became possible to determine for each matrix its “orthogonal” projection (by minimum distance) on the primary geodesic, producing both a value for its distance along the geodesic and its distance from the geodesic. A secondary geodesic was determined between the matrix which had the greatest distance from the initial geodesic and its projection on the primary geodesic. The method of the first analysis was then repeated, producing a value for the distance of each matrix along the secondary geodesic, and its distance from the geodesic. Visualization and interpretation of the results are provided by plotting the four distances obtained as trajectories along the age axis. The age at which the greatest distance from the primary geodesic occurred was also obtained. These values were calculated separately for males and females, and for the anterior and posterior blocks. It should be noted that in this particular analysis the initial and final matrices were set arbitrarily as being those of the initial and final ages; a more sophisticated analysis would use some algorithmic procedure or reference to the MDS analysis to determine their locations.

Bootstrapping was used to evaluate the robustness of the results, as explained subsequently for each type of analysis in Section 3.3. Bootstrap samples were constructed by random selection with replacement of observations, not of subjects. Each observation received a value of 0, 1, 2, … max number of times selected, and then when the weighted covariance calculations were performed, each weight was multiplied by the value received in the randomization procedure. One thousand bootstrap samples were created.

## 3 Results

### 3.1 Means and Variances of bipolar coherence

Means and variances increased from ages 12 to 20 and then remained relatively stable. Females initially had larger means than males, particularly in the anterior region, but sex differences for these measures were relatively small after age 18, differing between 5 and 10% of their mean value. Initially (ages 12-14) male and female covariances were similar; male covariances increased with age throughout the age span, but female covariances increased only to age 17 and were relatively stable thereafter.

To understand relation between rates of change and means, the zscores of the time series of means are shown in Figure 4. The patterns of development are clearly independent of the means.

**Figure 4:**
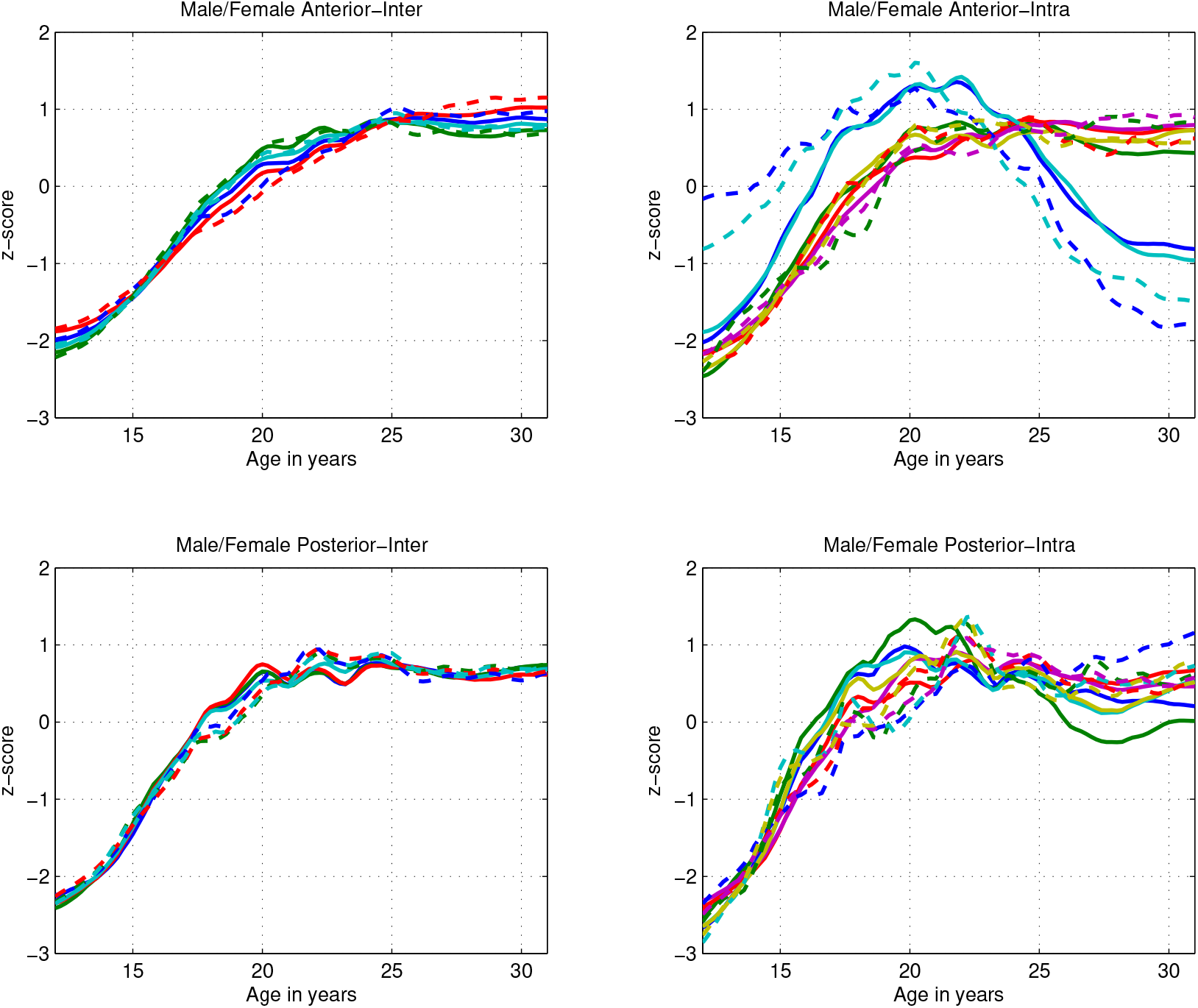
Z-scores of anterior and posterior high alpha band trajectories of the 27 bipolar coherence pairs used in this study. Coherence pairs were divided into 2 similar groups for each region: an inter-hemispheric group of 4 coherence pairs, and an intra-hemispheric and midline group of 6 coherence pairs. See Section 6.1 for complete lists. Male trajectories are represented by solid lines and female trajectories by dashed lines. The blue and cyan lines in the upper right panel are F7-C7 – F3-C3 and F7-C7 – FZ-CZ. Intrahemispheric pairs F7-F3 – T7-C3 and F7-F3 – P7-P4 also show anomalous trajectories not illustrated here.

**Figure 5:**
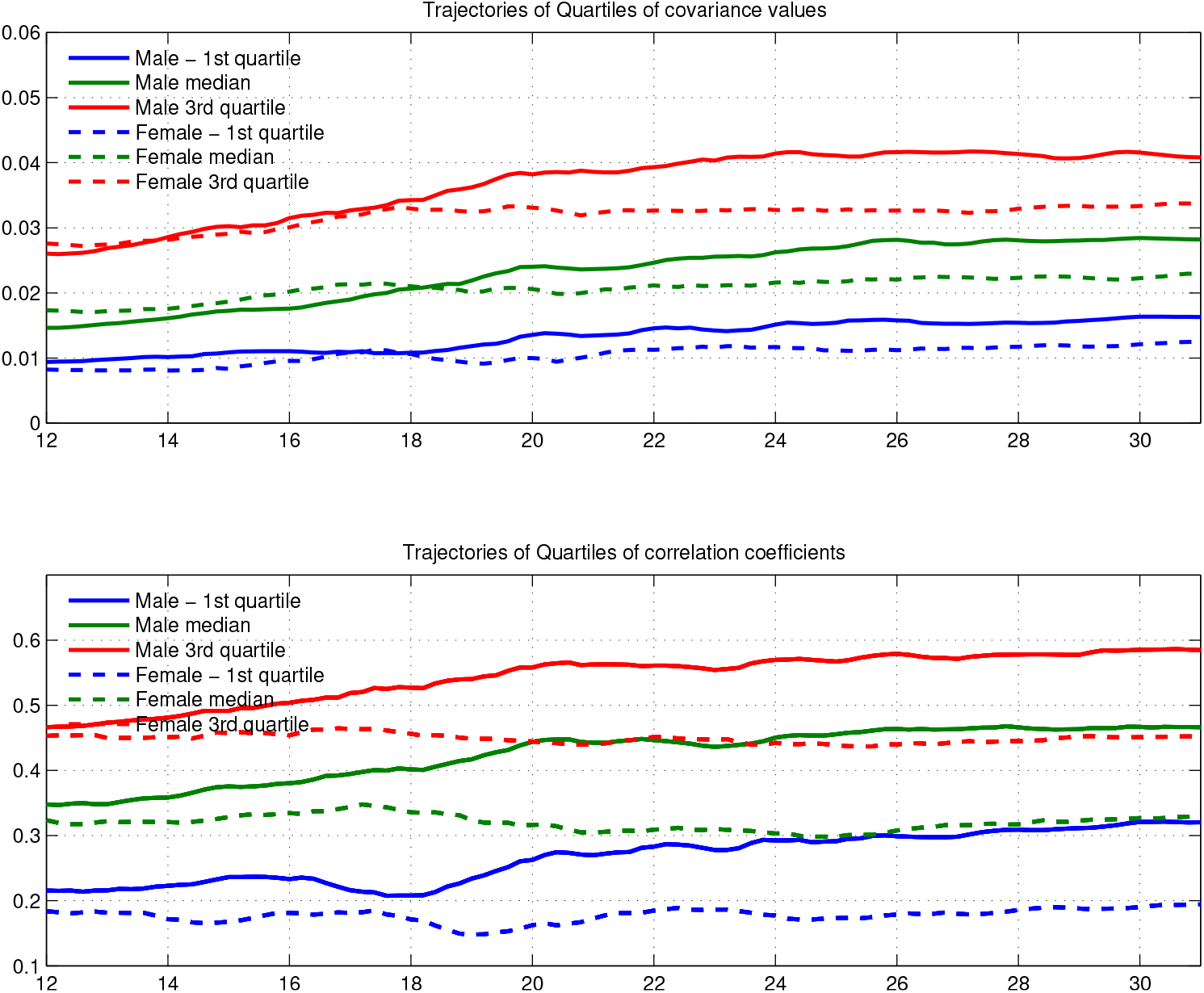
Upper Panel: Trajectories of the quartiles of the covariance values. Lower Panel: Trajectories of the quartiles of the correlation values.

### 3.2 Covariance analysis

#### Covariance and Correlation Trajectories of bipolar coherence

Quartiles of all male and female covariance and correlation trajectories are show to indicate general sex specific characteristics in the subsequent plot. Since there are 351 pairwise covariance/correlation values, a plot of all of their trajectories together would not be particularly informative. The linear order of the covariances is remarkably stable over age; the correlation of the covariances between different ages was greater than .9 over all pairs of ages. Since the female correlations are stable over the age range, the increase in female covariance driven only by an increase in variance, which in females is proportional to the mean.

#### 3.2.1 Tensorial Analysis

To provide an initial understanding of the relation between the coherence trajectories and the information provided by determining the geodesic distances between covariance matrices, a plot of the z-score of the geodesic distance between the initial (age 12) covariance matrix of the entire set of 27 coherence pairs and each successive covariance matrix was superimposed on the plot of the z-scores of the coherence trajectories (shown in Figure 4) is shown in Figure 6. The identity of temporal structure between the coherence trajectories and the geodesic distance trajectory suggests a direct connection between processes affecting development and the observed coherence values.

**Figure 6:**
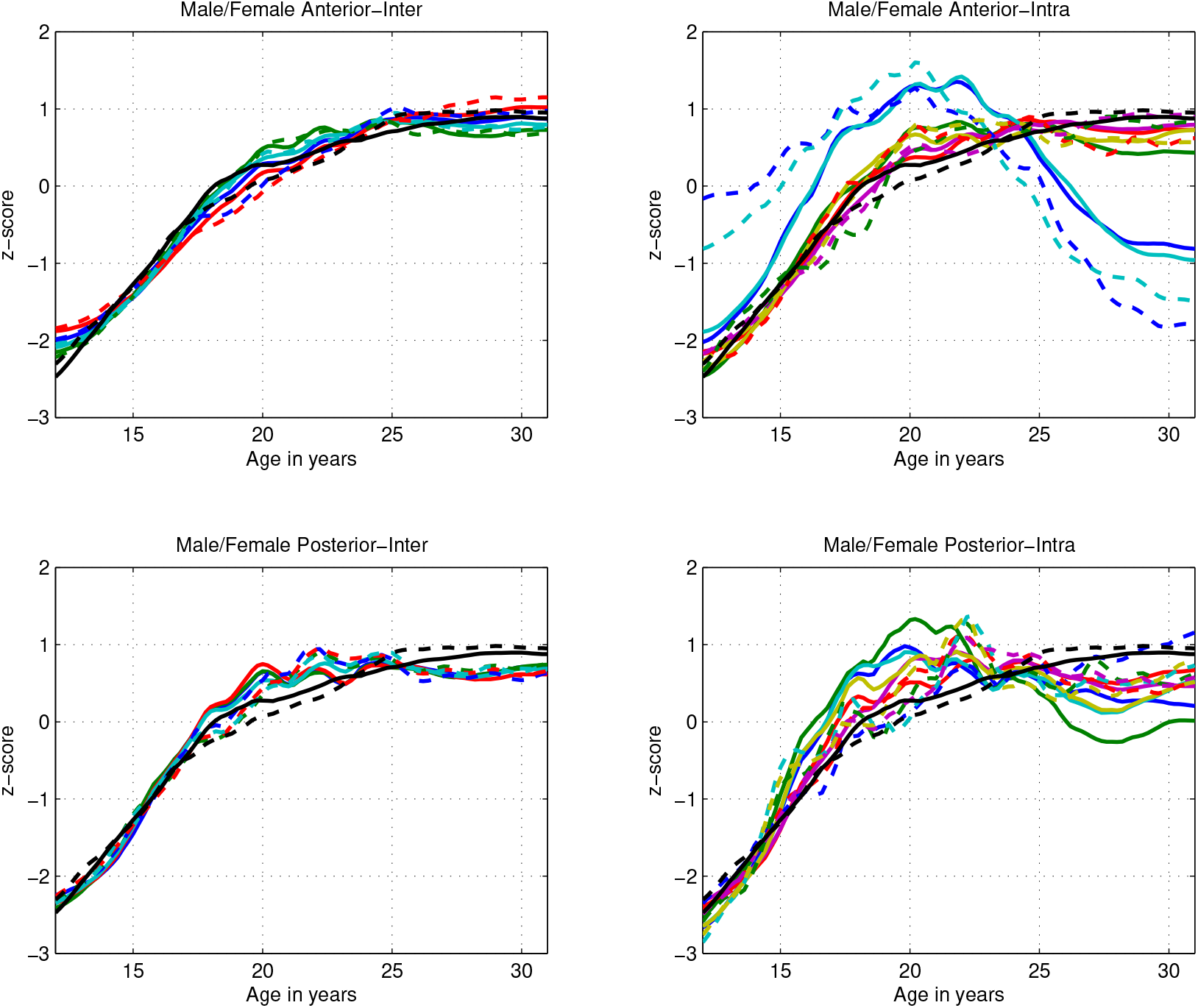
The z-score of the geodesic distance between the initial (age 12) covariance matrix of the entire set of 27 coherence pairs and each successive covariance matrix was superimposed on the Z-scores of anterior and posterior high alpha band trajectories of the 27 bipolar coherence pairs used in this study. Coherence pairs were divided into 2 similar groups for each region: an inter-hemispheric group of 4 coherence pairs, and an intra-hemispheric and midline group of 6 coherence pairs. See section 6.1 for complete lists. Male trajectories are represented by solid lines and female trajectories by dashed lines. The blue and cyan lines in the upper right panel are F7-C7 – F3-C3 and F7-C7 – FZ-CZ. Intrahemispheric pairs F7-F3 – T7-C3 and F7-F3 – P7-P4 also show anomalous trajectories not illustrated here. Geodesic distance trajectories are represented by black lines.

##### Multidimensional scaling

Results of the MDS calculations are shown in Figure 7, with the entire 27 coherence pair result shown first, with between sex comparisons. A comparative anterior/posterior and male/female set of MDS plots follows in Figure 8. In each case the trajectories for each axis are plotted against age; it is easier to interpret the trajectories in that way rather than as a space curve in 3 dimensions. Because the MDS attempts to represent the distances in its input, the unit of the y axes is the Riemannian distance between between the covariance matrices. The assumption in plotting the trajectories in a comparative manner is that the same axis (by algorithmic order) for each of the sets of covariance matrices represents the same aspect of the developmental process by temporal structure. If the shape of the trajectories along the same axis for different sets of covariance matrices were radically different, the interpretation of the plots would be different from that offered below. Note that the distance between the initial and final matrix in the MDS space is about 2.6 units; this is about a quarter of the estimate of the diameter of sets of comparable covariance matrices discussed in Section 2.3.1.

**Figure 7:**
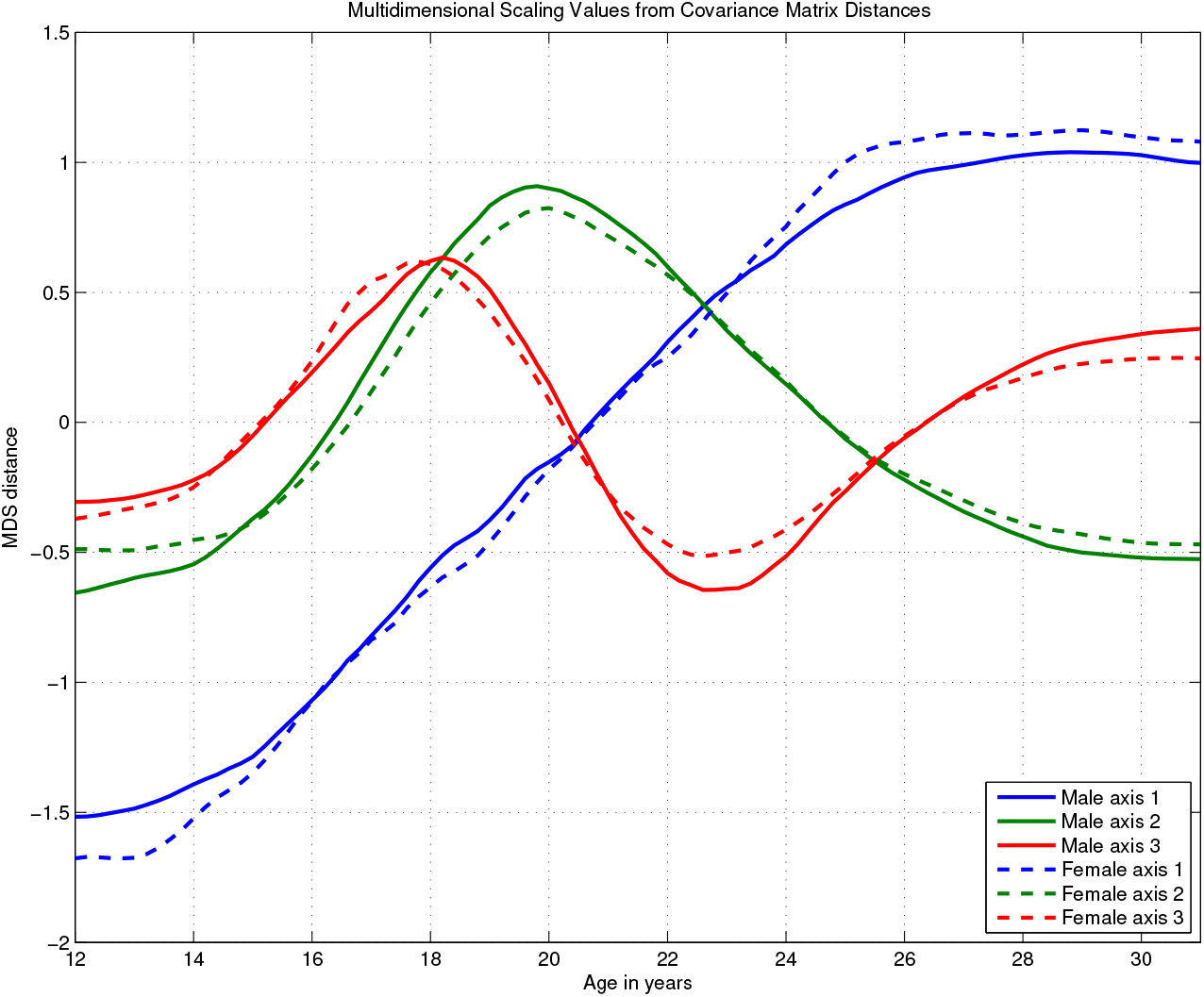
MDS trajectories for the complete covariance matrices.

**Figure 8:**
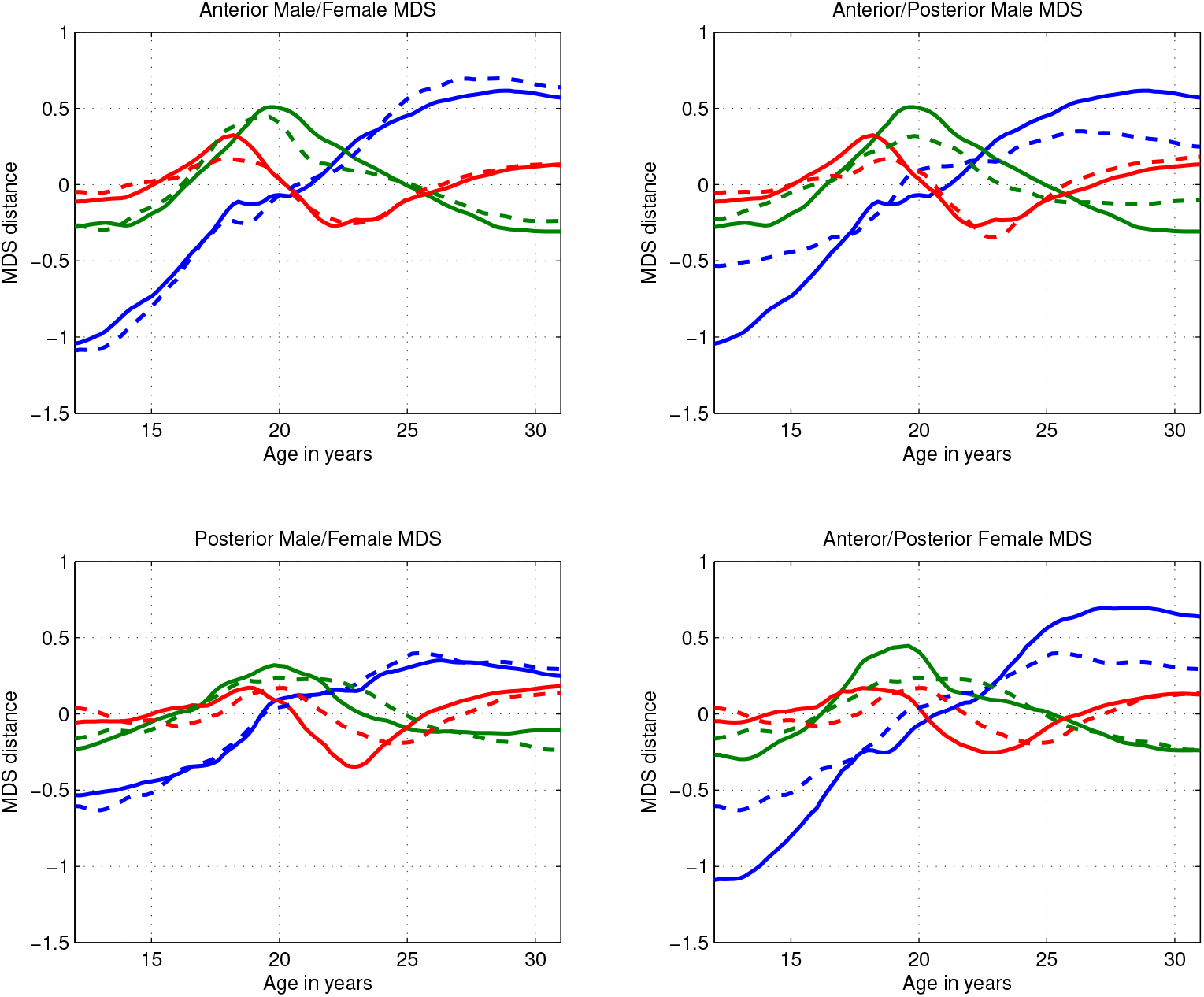
MDS trajectories for region/sex comparisons. Upper left: Anterior Male-Female comparison. Male lines are solid, Female dashed. Upper right: Male Anteror-Posterior comparison. Anterior lines are solid, Posterior dashed. Lower left: Posterior Male-Female comparison. Male lines are solid, Female dashed. Lower right: Female Anteror-Posterior comparison. Anterior lines are solid, Posterior dashed.

Given the shapes of the trajectories for each of the three axes, a plausible interpretation is that the trajectory along axis 1 (blue lines) represents a linear progression from an initial to final stable condition. The trajectory along axis 2 (green lines) represents an excursion from linearity. The trajectory along axis 3 (red lines) represents the asymmetry of ascending and descending limbs of the excursion. Note that final values along axis 2 are very similar to the initial values, while this is not the case for the values along axis 3. This indicates that the departure from linearity has both transient (axis 2) and lasting (axis 3) effects on the ultimate outcome of the development. This accords well with our initial interpretive framework.

What is striking about Figure 7 is the similarity of the male and female trajectories. In this analysis, the male trajectories are based solely on the distances between male covariance matrices, the female trajectories solely on the distances between female covariance matrices; no direct male-female distances are shown here. The calculated male-female distances are shown in Figure 9 below.

**Figure 9:**
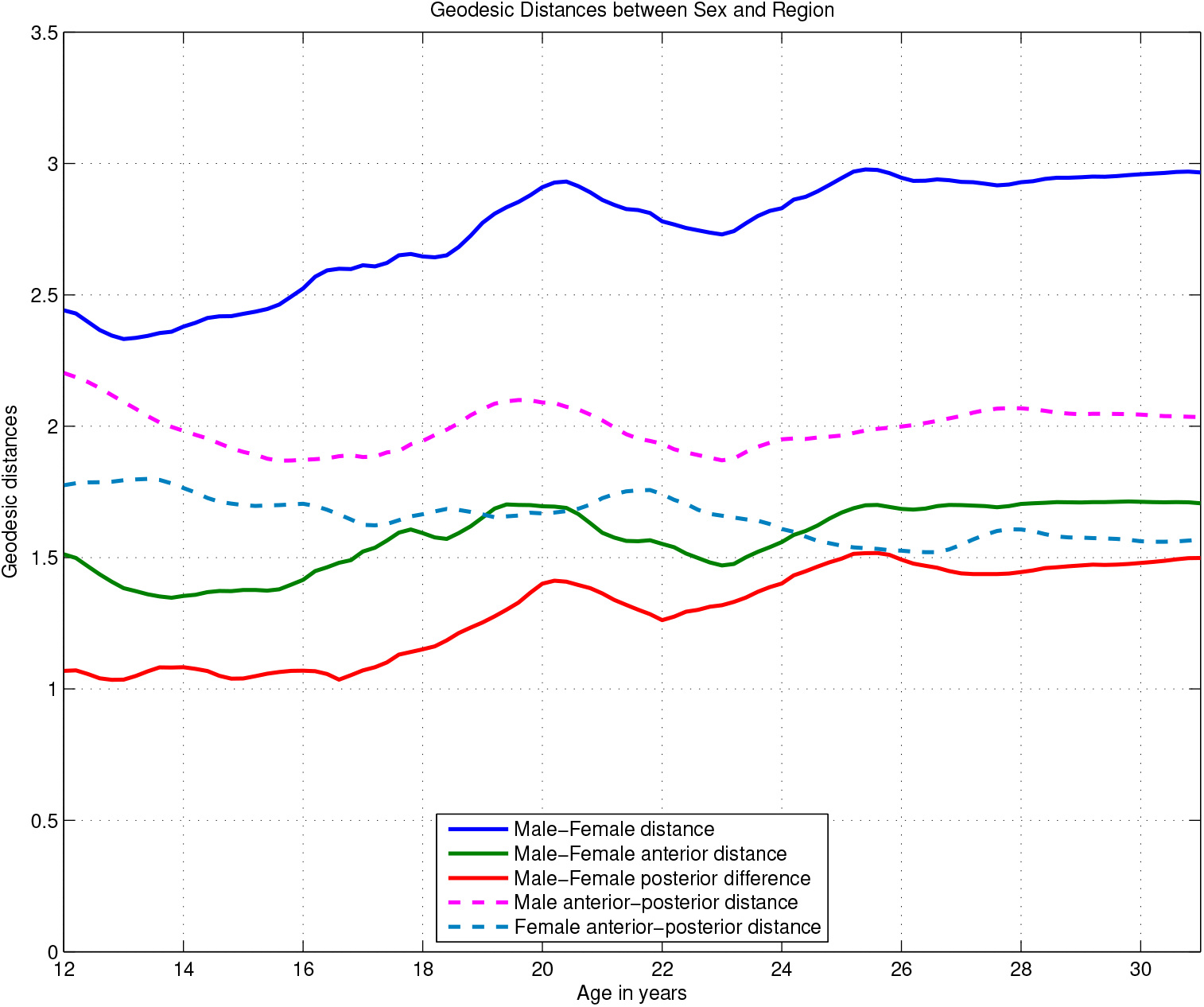
Trajectories of Riemannian distances between males and females by region and between regions within sexes.

In Figure 8 which illustrates sex differences by region (left column) and regional differences by sex (right column), it is clear that sex differences by region are small compared to regional differences in each sex. In the right column, the difference between the regions is most significant along the first axis, which represents the initial to final progression, with the anterior progressions strikingly larger than the than the posterior progressions for both sexes. In the left column, this is reflected in the that the range of values for the y axis is considerable less in the posterior region than in the anterior region. This is the result of the factor that there is far less age-related variability in the covariances between posterior coherence pairs than between anterior coherence pairs.

As mentioned above, these MDS trajectories are based upon within sex and within region distances. When distances between sexes are examined directly, as shown in the subsequent plot. they show relatively little variability with age, and are only weakly related to the differences between sexes and regions shown in the previous plot.

##### Riemannian Distances between sexes and regions

The distances between the matrices for males and females are shown in Figure 9, they are about the same size as the differences between the initial and final matrices in the each of the sexes individually. This, taken with the MDS results shown above is by far the most important result of the analysis: the precise parallelism of male and female MDS trajectories with the consistent difference in distance between the corresponding male and female covariance ages. The fact that male covariance values are larger than the female covariance values after age 18 (see Figure 5) cannot be a complete explanation for this.

##### Geodesic Linearity Analysis

In contrast with the MDS plots, the GLA plots only provide two axes, although the plots of the distance from the trajectories does offer additional information. In the plots, the solid blue line represents the linear trend of the projection of the covariance matrices along the primary geodesic, and the dashed blue line how far away these matrices are from the primary geodesic. The solid green line represents the linear trend of the projection of covariance matrices along the secondary geodesic, and the dashed green line how far away these matrices are from that geodesic. The distances on the y-axis are those of the non-Euclidean space of the covariance matrices. In the comprehensive plot, The distance of the excursion from the linear progression is about half of the distance from the initial to the final state. Note that the age at maximum excursion from linearity is the same for males and females, and is the same as the peak in the trajectory along axis 2 in the MDS analysis. The similarity between male and female trajectories is parallel to the similarity in the MDS plots. As suggested in Section 2, the plot might have been more informative had the ages defining the geodesic been closer to the initial and final points of the ascending part of MDS trajectory in axis 1. (By definition, the age at which the distance of the observed matrix from the linear geodesic is greatest is identical to the age at which the distance from the secondary geodesic is zero. Also by definition the distances along the secondary geodesic must be zero at the points chosen as the determinants of the primary geodesic.)

As in the MDS plots regional differences are larger than sex differences. Particularly noticeable is the greater distance between the initial and final matrices for the anterior region, as well as the earlier peak of the excursion from linearity compared to the posterior region. In contrast to the MDS plots, where the linear progression emerged directly from the Euclidean representation of the data, the linear progression was assumed in the analysis procedure, but if there had been a greater deviation from linearity than that observed, the solid blue line in the plots would not be so close to linear.

### 3.3 Bootstrap procedures and results

#### Bootstrap procedures and results

Bootstrap evaluation for the MDS trajectories shown in Figures 7 and 8 was conducted by calculating the correlation coefficients between the observed male and female trajectories along the corresponding axes and exhibited on the left side of Table1 under “Observed Correlations”; then for each bootstrap sample the same correlations were calculated and their mean over the 1000 bootstrap samples was determined and shown on the right side of Table 1 under “Bootstrap Mean Correlations”. Additionally, the correlation coefficients between the observed trajectories and each of the corresponding bootstrap sample trajectories was calculated and the means are shown in Table 2 for each sex.

**Table 1:** Observed and mean bootstrap correlation coefficients between males and females for each of the 3 MDS axes

| Observed Correlations |  |  |  | Bootstrap Mean Correlations |  |  |  |
| --- | --- | --- | --- | --- | --- | --- | --- |
| Region |  |  |  | Region |  |  |  |
|  | Anterior | Posterior | Entire |  | Anterior | Posterior | Entire |
| Axis |  |  |  |  |  |  |  |
| 1 | 0.9968 | 0.9951 | 0.9988 |  | 0.9828 | 0.9628 | 0.9937 |
| 2 | 0.9606 | 0.8932 | 0.9963 |  | 0.9302 | 0.8417 | 0.9793 |
| 3 | 0.9532 | 0.6539 | 0.9842 |  | 0.8734 | 0.7036 | 0.9648 |

**Table 2:** Mean correlation coefficients for correlation between observed and bootstrap values for males and females for each of the 3 MDS axes

| Male Mean Correlations |  |  |  | Female Mean Correlations |  |  |  |
| --- | --- | --- | --- | --- | --- | --- | --- |
| Region |  |  |  | Region |  |  |  |
|  | Anterior | Posterior | Entire |  | Anterior | Posterior | Entire |
| Axis |  |  |  |  |  |  |  |
| 1 | 0.9904 | 0.9766 | 0.9947 |  | 0.9931 | 0.9813 | 0.9965 |
| 2 | 0.9686 | 0.8981 | 0.9892 |  | 0.9602 | 0.9340 | 0.9909 |
| 3 | 0.9521 | 0.8618 | 0.9856 |  | 0.9016 | 0.5107 | 0.9869 |

Corresponding results for the GLA trajectories, shown in Figures 10 and 11, are in in tables 3 and 4.

**Table 3:**
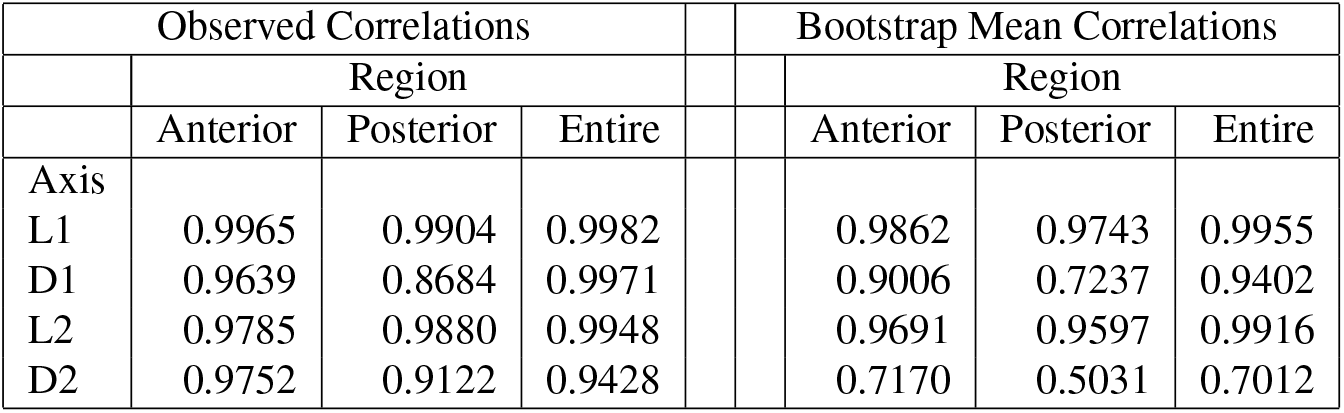
Observed and mean bootstrap correlation coefficients between males and females for each of the 4 Geodesic linearity trajectories

| Observed Correlations |  |  |  | Bootstrap Mean Correlations |  |  |  |
| --- | --- | --- | --- | --- | --- | --- | --- |
| Region |  |  |  | Region |  |  |  |
|  | Anterior | Posterior | Entire |  | Anterior | Posterior | Entire |
| Axis |  |  |  |  |  |  |  |
| L1 | 0.9965 | 0.9904 | 0.9982 |  | 0.9862 | 0.9743 | 0.9955 |
| D1 | 0.9639 | 0.8684 | 0.9971 |  | 0.9006 | 0.7237 | 0.9402 |
| L2 | 0.9785 | 0.9880 | 0.9948 |  | 0.9691 | 0.9597 | 0.9916 |
| D2 | 0.9752 | 0.9122 | 0.9428 |  | 0.7170 | 0.5031 | 0.7012 |

**Table 4:** Mean correlation coefficients for correlation between observed and bootstrap values for males and females for each of the 4 Geodesic linearity trajectories

| Male Mean Correlations |  |  |  |  | Female Mean Correlations |  |  |  |
| --- | --- | --- | --- | --- | --- | --- | --- | --- |
|  | Region |  |  |  |  | Region |  |  |
|  | Anterior | Posterior | Entire |  |  | Anterior | Posterior | Entire |
| Axis |  |  |  |  |  |  |  |  |
| L1 | 0.9936 | 0.9828 | 0.9975 |  |  | 0.9945 | 0.9884 | 0.9980 |
| D1 | 0.9512 | 0.3675 | 0.9710 |  |  | 0.9570 | 0.7731 | 0.9663 |
| L2 | 0.9871 | 0.9679 | 0.9962 |  |  | 0.9816 | 0.9747 | 0.9953 |
| D2 | 0.8365 | 0.1788 | 0.8069 |  |  | 0.8617 | 0.3462 | 0.8324 |

**Figure 10:**
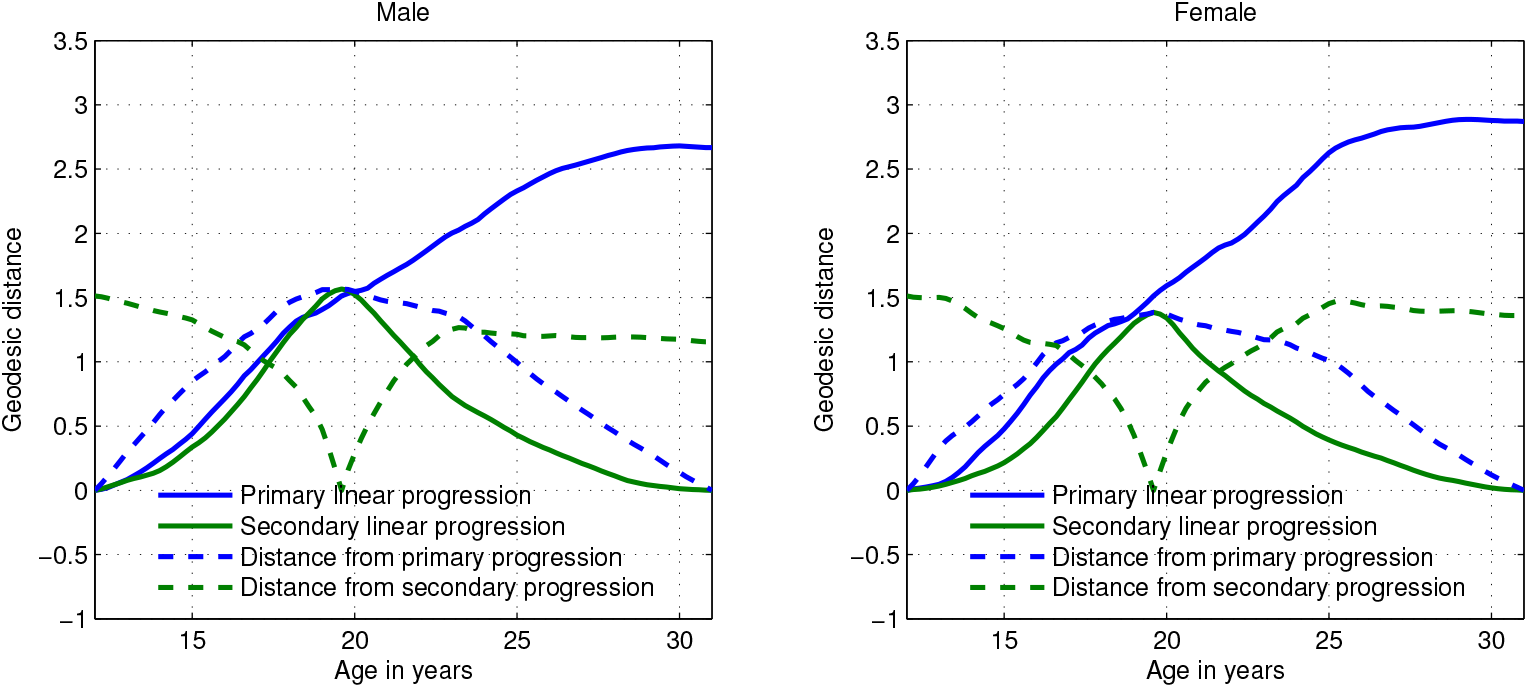
Geodesic linearity analysis of entire system.

**Figure 11:**
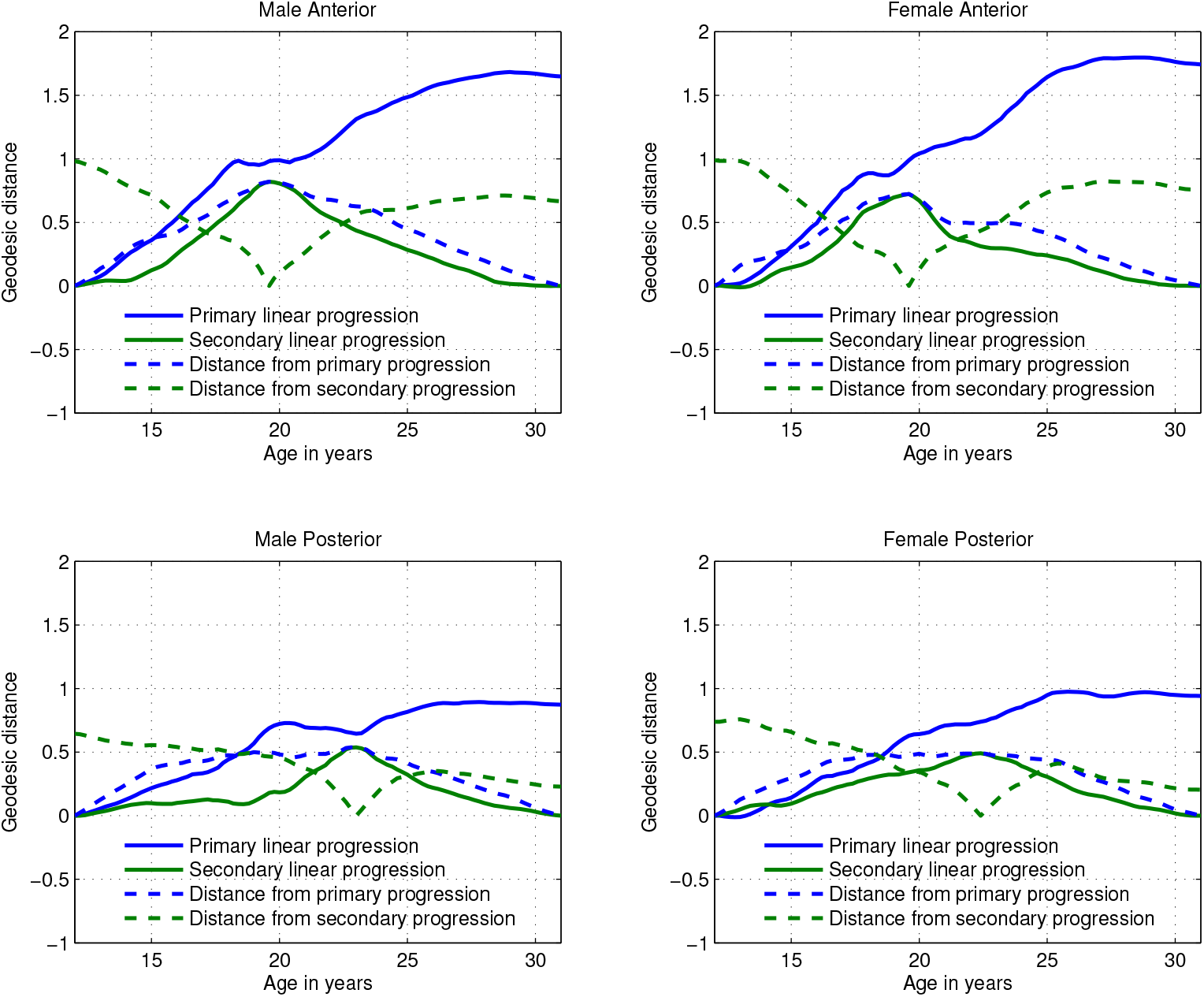
Geodesic linearity analysis of Anterior and Posterior subsystems.

#### Bootstrap results observations

For the MDS analysis, only the posterior female trajectory is not a robust result, with the correlation of the observed results with the bootstrap results significantly lower than any of the other as shown in Table 2. For the GLA analysis, the D “axes”, the distance from the primary and secondary geodesics, is less robust than the distances along the geodesics, particularly for the posterior region.

## 4 Discussion

### 4.1 Overview of results

Results are consistent with the following model:

1. The increase in coherence from ages 12 to 20 is driven by a relatively large number of age and topographically specific factors:
  - Short term fluctuations of covariance trajectories (not analyzed here) resulting from age-varying (between individuals) onset and cessation of influence of individual factors which produce respectively intervals of decorrelation and recorrelation between different coherence pairs.
  - Continued increase in geodesic distance from initial covariance matrix of subsequent covariance matrices is consistent with the increases in coherence.
2. After age 20, new factors diminish in number, and cessation of previously active factors occur:
  - Coherence values remain stable, but correlations in males increase slightly, while female correlations have a mixed pattern of increase and decrease.
  - There is less change in covariance matrices, particularly after age 25.

Analysis of individual coherence trajectories by themselves cannot determine the degree of relatedness between the parallel trajectories which constitute the bulk of the results. Clearly the tensorial analysis using MDS suggests a more complex developmental process than the bipartite age divisions given by the description of the means and variances of the coherence values by themselves. The MDS results suggest an initial stage of concerted development from age 14 to 18, a complex reorganization from age 18 to 25, and a period of stabilization after age 25. The tensorial analysis using GLA indicates age 18 as an important transition point. Some aspects of the GLA plots suggest a change in the relation between anterior and posterior subsystems at that transition.

### 4.2 Research Directions

Preliminary work on the fine structure of the covariance matrices and of individual pairwise trajectories and their derivatives in this and other similar data sets suggest that these measures will be able to identify specific topographies with the elements of the overall trajectories shown by the MDS and GLA results. Additionally, more coherence pairs should be studied to understand the left-right relations with the detail provided here to the anterior-posterior relations.

Our discussion of genetic associations of the individual neurophysiological measures found in Chorlian et al. (2017); Meyers et al. (2019, 2021); Chorlian et al. (2023) and inspired by Stead et al. (2006) and of the utilization of the covariance of association trajectories (via sparse PCA) in Chorlian et al. (2023) suggests that some way of bringing these elements together with analysis presented here may provide concrete evidence for some of the ideas quoted from Mitteroecker and Bookstein (2009) in our introductory section.

## 5 Acknowledgments

The Collaborative Study on the Genetics of Alcoholism (COGA), Principal Investigators B. Porjesz, V. Hesselbrock, T. Foroud; Scientific Director, A. Agrawal; Translational Director, D. Dick, includes ten different centers: University of Connecticut (V. Hesselbrock); Indiana University (H.J. Edenberg, T. Foroud, Y. Liu, M.H. Plawecki); University of Iowa Carver College of Medicine (S. Kuperman, J. Kramer); SUNY Downstate Health Sciences University (B. Porjesz, J. Meyers, C. Kamarajan, A. Pandey); Washington University in St. Louis (L. Bierut, J. Rice, K. Bucholz, A. Agrawal); University of California at San Diego (M. Schuckit); Rutgers University (J. Tischfield, D. Dick, R. Hart, J. Salvatore); The Children’s Hospital of Philadelphia, University of Pennsylvania (L. Almasy); Icahn School of Medicine at Mount Sinai (A. Goate, P. Slesinger); and Howard University (D. Scott). Other COGA collaborators include: L. Bauer (University of Connecticut); J. Nurnberger Jr., L. Wetherill, X., Xuei, D. Lai, S. O’Connor, (Indiana University); G. Chan (University of Iowa; University of Connecticut); D.B. Chorlian, J. Zhang, P. Barr, S. Kinreich, G. Pandey (SUNY Downstate); N. Mullins (Icahn School of Medicine at Mount Sinai); A. Anokhin, S. Hartz, E. Johnson, V. McCutcheon, S. Saccone (Washington University); J. Moore, F. Aliev, Z. Pang, S. Kuo (Rutgers University); A. Merikangas (The Children’s Hospital of Philadelphia and University of Pennsylvania); H. Chin and A. Parsian are the NIAAA Staff Collaborators.

We continue to be inspired by our memories of Henri Begleiter and Theodore Reich, founding PI and Co-PI of COGA, and also owe a debt of gratitude to other past organizers of COGA, including TingKai Li, P. Michael Conneally, Raymond Crowe, and Wendy Reich, for their critical contributions. This national collaborative study is supported by NIH Grant U10AA008401 from the National Institute on Alcohol Abuse and Alcoholism (NIAAA) and the National Institute on Drug Abuse (NIDA).

## Historical Note

The results presented here were the topic of a presentation given in 2020 to the COGA Brain Function group. No further analysis of this data set has been done since then. A copy of the 2020 presentation can be obtained from the first author.

## 6 Supplemental Material

### 6.1 Block structure of coherence pairs

Sagittal Bipolar:

Anterior Interhemispheric (Frontal-Central)

1 F8-T8 – F7-T7

2 F4-C4 – F3-C3

3 F3-C3 – F8-T8

4 F4-C4 – F7-T7

Anterior Midline and Intrahemispheric (Frontal-Central)

5 F3-C3 – F7-T7

6 F4-C4 – F8-T8

7 FZ-CZ – F7-T7

8 FZ-CZ – F3-C3

9 FZ-CZ – F8-T8

10 FZ-CZ – F4-C4

Posterior Interhemispheric (Central - Parietal)

11 T8-P8 – T7-P7

12 C4-P4 – C3-P3

13 C3-P3 – T8-P8

14 C4-P4 – T7-P7

Posterior Midline and Intrahemispheric (Central - Parietal)

15 C3-P3 – T7-P7

16 C4-P4 – T8-P8

17 T7-P7 – CZ-PZ

18 C3-P3 – CZ-PZ

19 T8-P8 – CZ-PZ

20 C4-P4 – CZ-PZ

Parietal – Occipital

21 P4-O2 – P3-O1

Lateral Bipolar:

Left Intrahemispheric

22 T7-C3 – F7-F3

23 P7-P3 – F7-F3

24 P7-P3 – T7-C3

Right Intrahemispheric

25 T8-C4 – F8-F4

26 P8-P4 – F8-F4

27 P8-P4 – T8-C4

### 6.2 Mathematical Formulas for Log-Euclidean computation

Given X is PDS and [U, S, V] = svd(X) such that U * S * V^′^ = X, we have

log(X) = U * log(S) * V^′^

exp(X) = U * exp(S) * V^′^

The distance between two PDS matrices X_1_ and X_2_ is then defined as

dist(X_1_, X_2_) = trace((log(X_1_) − log(X_2_))^2^)^1*/*2^

and the geodesic between X_1_ and X_2_ is

G(t) = exp((1 − t) log(X_1_) + t log(X_2_)) as described in Arsigny et al. (2006).

Matlab functions using the above definitions were used for all of the covariance matrix analysis in this paper.

